# Detection for suspected Genetically Modified Maize and Soybean crops in the selcted places of Ethiopia using PCR machine

**DOI:** 10.1101/2019.12.12.874974

**Authors:** Adugnaw Admas, Smegnew Melese, Amare Genetu

## Abstract

Detection of genetically modified organisms (GMOs) in crops is an important issue for all the subjects involved in seed quality control and customer’s right. Due to the increasing number of GMOs research and development activity in the globe during the past few years, it has become necessary to screen and regulate highely practiced GMO crops. In this study, following the extraction of genomic DNA from agriculture farming land and commercial areas collected samples that suspected for GMOs such as Soybean and maize crops, the recombinant DNA target sequences were targeted to detect for transgene such as CaMV35s gene using thermal cycler Polymerase Chain Reaction (PCR). In addition, for positive control taxone specfic *Invertase* and *Lectine* gene of maize and soybean have done. Based on the gel electrophoresis results, *Invertase and Lectine* gene events were detected in all maize and soybean samples respectively. How ever, fortunately in all collected samples of maize and soybean in different places of the countery trans-genes were not detected. Finally, this reserch result concludes that in the studied area soybean and maize crops are GMO free.

## 1. Introduction

Now a day there has been exponential growth in the field of modern biotechnoloy or genetic engineering for trait improvements during the past 20 years. This result brought to genetically modify agricultural crops for potential improvement in crop quality and productivity. Plants produced by transgenic technology or genetic engineering are called genetically modified organism (GMO). In recent years surplus number of crop plant species have been genetically modified to attain traits such as biological defense against diseases and insects, thus reducing the need for chemical pesticides and introducing genetic traits that enable crops to better withstand drought, temprature, saline conditions and in addition, genetic modification has been used to improve various characteristics such as color, flavor, texture, nutrient value and shelf life of foods [1-7].

Many genetically modified crops have been approved worldwide since mid-1990s and the global area of these crops has expanded yearly since their commercialization. In 2011, 160 million hectares were cultivated with genetically modified crops mainly soybean, maize, cotton and canola [8]. A total of 26 countries, 19 developing and 7 industrial countries planted biotech crops in 2016. The top ten countries, each of which grew over 170 million hectares in 2016, is led by the USA which grew 72.9 million hectares (39% of global total, similar to 2015), Brazil with 49.1 million hectares (27%), Argentina with 23.8 million hectares (13%), Canada with 11.1 million hectares (6%), India with 10.8 million hectares (6%), Paraguay with 3.6 million hectares (2%), Pakistan with 2.9 million hectares (2%), China with 2.8 million hectares (2%), South Africa with 2.7 million hectares (1%) and Uruguay with 1.3 million hectares (1%). An additional 16 countries grew a total of approximately 4.9 million hectares in 2016 [9]. Also, in Ethiopia Bt-Cotton is enviromentally realeased in may 2018 after amenmdments of biosafety regulation in august,2015 to use GMO for commerial and research purpose] Ethiopian Biosafety regulation, 2015] and other africa countery such as Burkine Faso, Sudan and Nigeria were allowed recentelly certain GMO crops for application.

At the moment, several transgenes have been approved worldwide for cultivation and consumption as food and feed, while consumer›s increasing awareness regarding food safety has created the need for more stringent regulations to control raw materials, import, export and distribution of GMOs [10, 11]. Therefore, the detection and identification of GMOs in food and seed have become important issues for all the subjects involved in food and import and export seed control [12, 13]. Qualitative testing may be used to discriminate between the authorized and unauthorized GM food to identify safe or potentially unsafe material [14, 15].

Although, several analytical methods have been proposed, among GMO detection methods generally involve specific DNA sequence detection by means of PCR techniques able to detect even small amounts of transgenes in raw materials and processed foods [16, 17]. Hence in this study, we have examined the suspected recombinant DNA of genetically modified maize and soybean in different agriculture farming land and commercial aeras of Ethiopia.

## 2. Materials and methods

### 2.1. Maize and soybean crops sample collection

In the Selected commercial and Agriculture farming areas of Ethiopa, specifically Areba Minich, Gondar, Bahrdar, Awashe sebate, AddisAbaba, Awi Zone at Ayehu mechanized agriculture farming land and Chagni soybean and maize crops sample were collected during the period Septmber 2018-June 2019.

### 2.2. Genomic DNA extraction of Soybean and Maize

Genomic DNA extraction was performed by transfering 1 g powder of the growended soybean and maize by high speed multi-functional crusher that can rotate 25000 RPM into a 50 ml falcon tube. In the flacon tube 30 ml CTAB Extraction buffer and 2.1 ml 20% SDS were added. By follwing this, the mixture was incubated at 65°C for 30 minutes. To separet the supernatant from pellet the sample was Centrifuged for 20 minutes at 3,000 × g, then after centrifugtion the deberis of the cell was exsited in the pellet phase but DNA material was existed in the supernatant phase, therefore, 20 ml supernatant was transferred to a new 50 ml falcon tube using a 5 ml pipette.In the transferd DNA material 5 ml of 5M KAC was added and incubated on ice for 30 minutes. Then, it was centrifuged for 20 minutes at 3,000 × g and 20 ml supernatant was transferred to a new 50 ml falcon tube using a 25 ml pipette, also added equal volume of isopropanol by follwing this incubate on ice for 5 minutes and Centrifuged for 20 minutes at 3,000 × g. Finally, the supernatant was discarded and the pellet was air-dried at 37°C until all isopropanol residues were evaporated then the pellet was completelly dissolved in 10 ml TE for PCR reaction and stored at -20 °C for futher use.

#### 2.2.1. Determination of purity of extracted DNA

The purity of extracted DNA were tested by 1.2 % Agarose gel electrophoresis.

**Figure 1.**
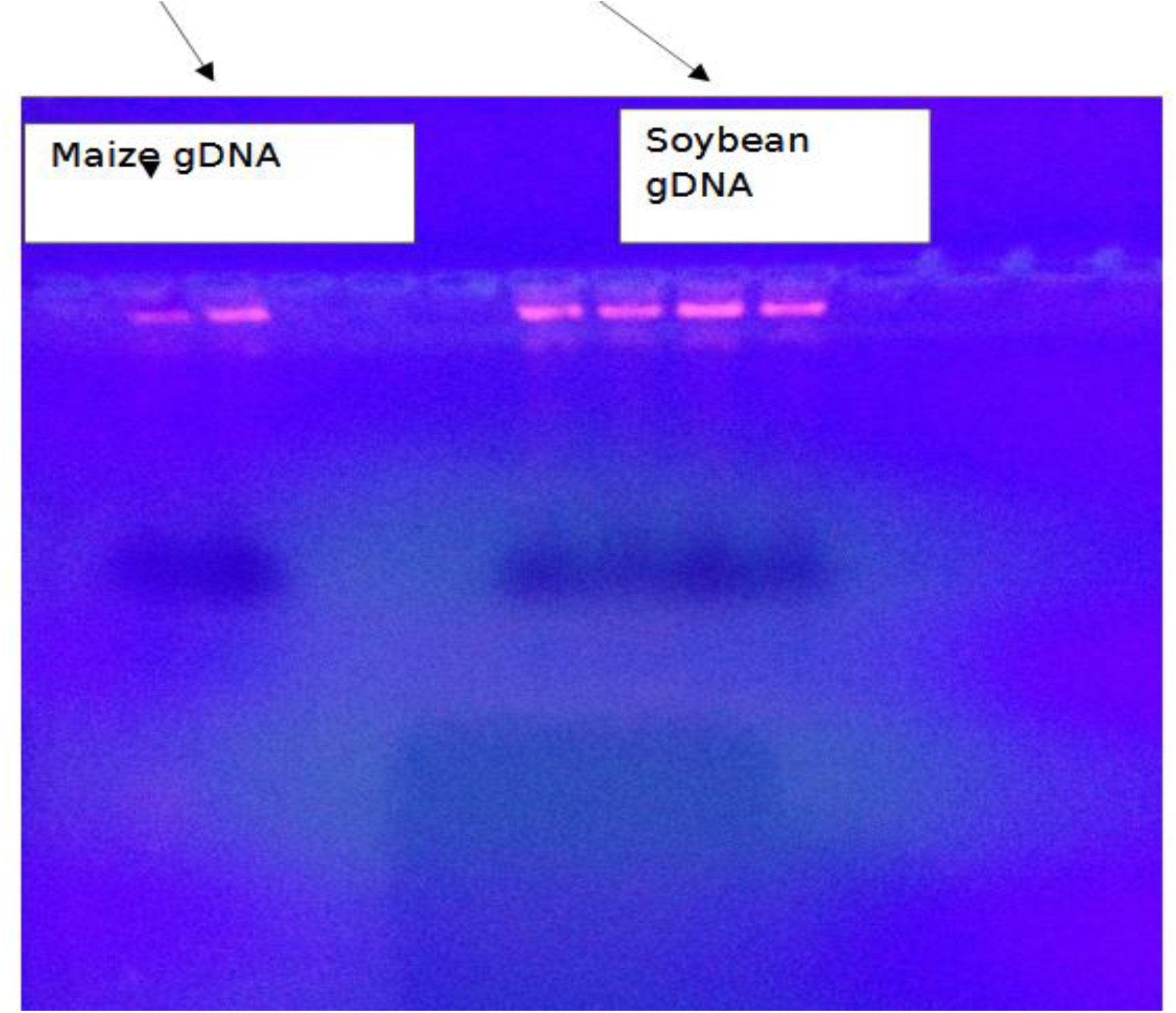
Extracted DNA from maize and Soybean.

### 2.3. DNA amplifications

PCR reactions were carried out in a total volume of 20 μL. Each reaction mixture contained 2 μL of 10x PCR buffer, 1.6 μL of 25 mM MgCl_2_, 0.2 μL of Taq DNA polymerase, 1.6 μL of 2.5 mM dNTPs, 4 μL of extracted DNA, 2 μL of 10 μM for all foreward and reverse Primer and 6.6 μL nuclease-free water were used for a reaction to detect the expected gene from DNA of maize and soybean. Thermal cycler conditions were as follows: pre incubation at 95°C for 5 min, 40 cycles consisting of dsDNA denaturation at 95°C for 20 sec, primer annealing for 40 sec at 57°C for extension at 72°C for 60 sec and final elongation at 72°C for 3 min.

Amplification PCR products of 5 μl of each and 3 μl loading dye were electrophoresed into 2 % agarose gel in 1x TBE buffer, stained with 5 μl ethidium bromide and visualized by gel doucumentation system with ultraviolet light and the amplification products were determined by using 3 μl of 100 bp DNA marker.

**Table 1.**
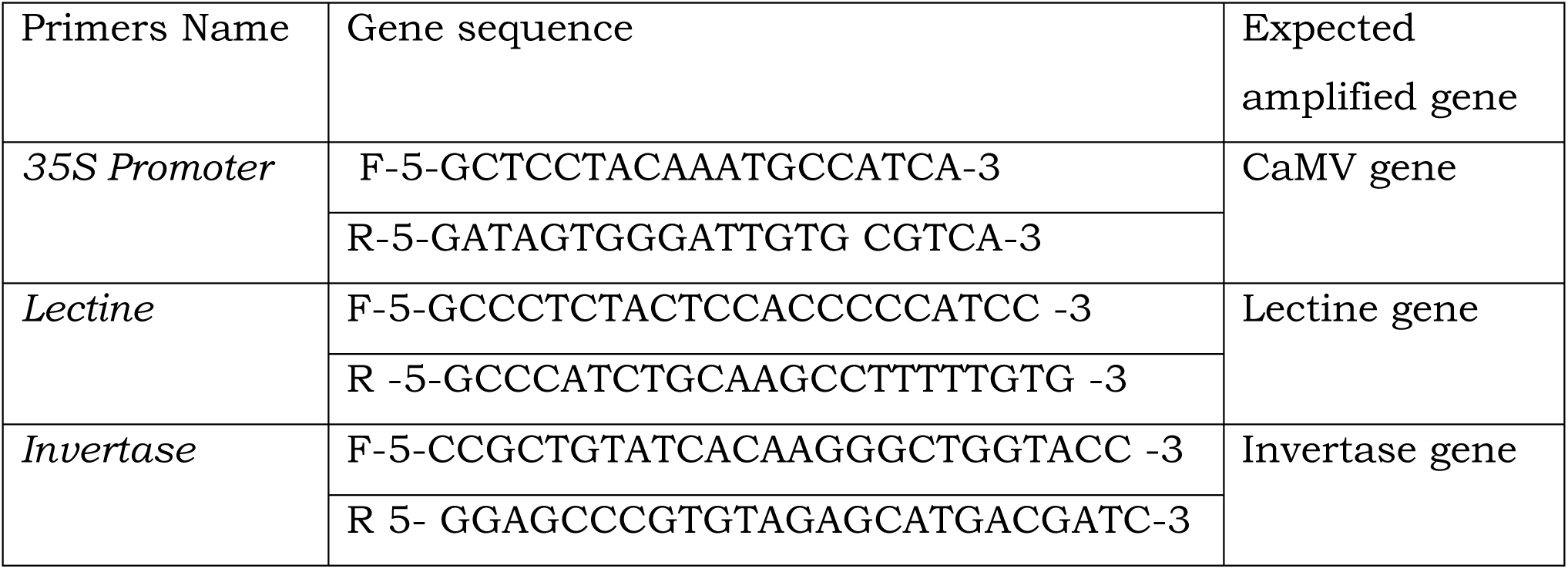
Primers were used for GMO detection and its sequence.

## 3. Result

Normal thermal cycler PCR-based method were used to detect the suspected crops for genetically modified maize and soyabean. PCR reactions were performed by using a 35S promoter-specific primer and the tareget plant host gene primer for *Invertase* and *Lectine* gene as positive control. The PCR products were analyzed in electrophorosis via 2% agarose gel.

The primer pair Lectine, specific for the single copy endogenous *lectin* gene in the collected samples of soybean were detected and its PCR product were 118 bp size as mentioned earlier by **Meyer *et al* [18]**. The other primer pair *Invetrase*, specific for the endogenous gene *Invertas*e was amplified to give a 226 bp amplicon **[19**]. This product was also detected in all collected sampls of maize crops. However, the desirable PCR products of 195 bp fragment of CaMV 35S promoter was not amplified in soybean and maize.

Therefore, these primers can be used for the routine screening and detection of specific gene in suspected GM soybean and maize. Most of the new analytical methods used for detection and quantification of transgenes in GM plants depend upon real time PCR assays, DNA-Chip technologies and using GMO screeing kit which are very expensive. However, the inexpensive thermal cycler and gene specfic primer method used in the present study would be more suitable for meeting regulatory obligations in developing countries.

One hybrids of GM cotton were approved in Ethiopia for commercialization in May 2018. However, the detection methods on different levels, such as qualitative, quantitative, national or international validated and non-validated, have been not developed to cheack really the commercialized Cotton is GMO or not, these detection methods and monitoring programmes of GM food/derivatives, import and export crops were not established in Ethiopia for a long time. But, time has come now to put in place the detection procedures, protocols and established a laboratory for GM planting material in different crops by Ethiopian Enviroment and Forest Research Institute.

**Figure 2.**
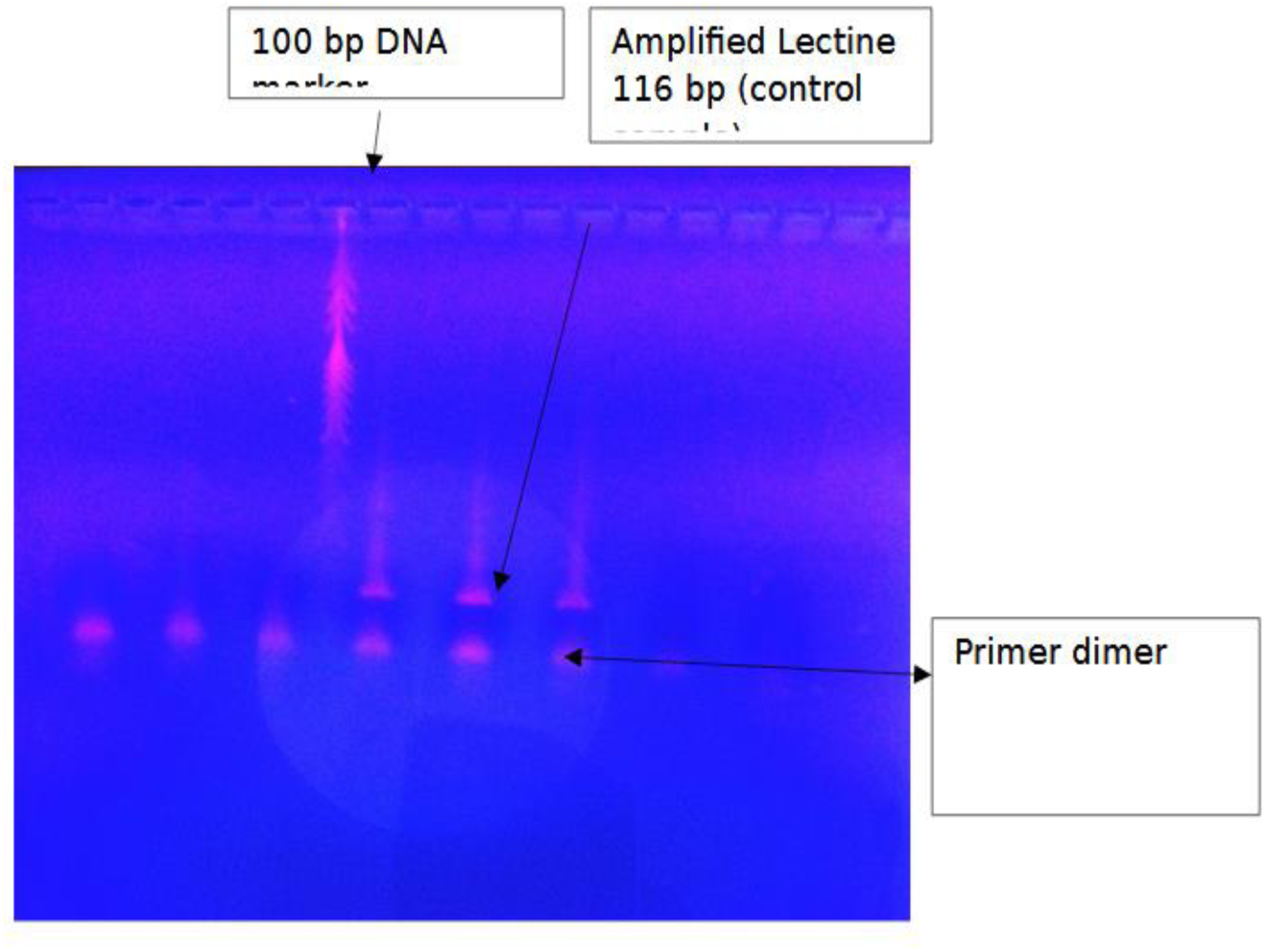
Amplified PCR products of soybean.

**Table 1:**
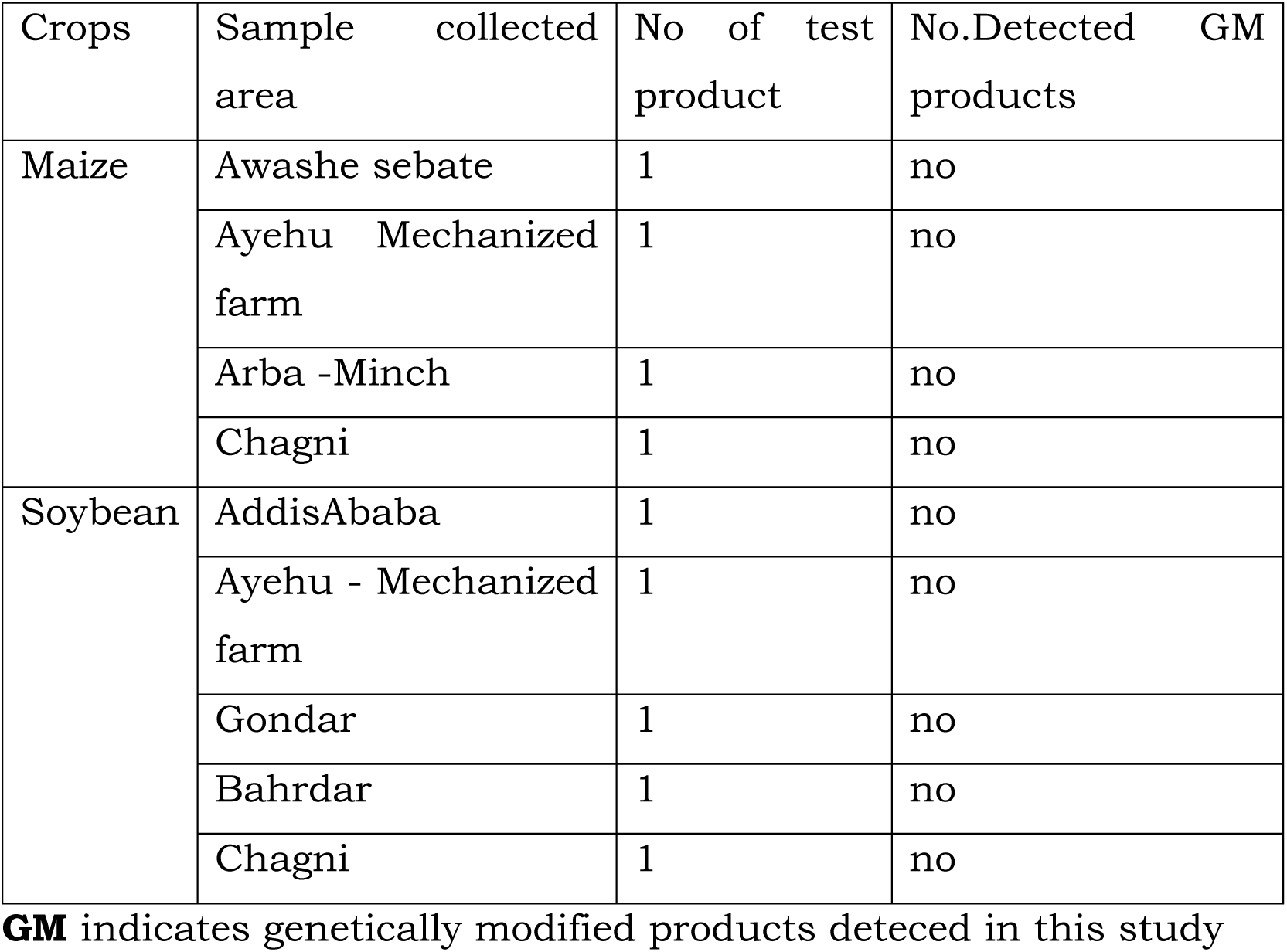
Areas of analyzed maize and soybean crops.

## 6. DISCUSSION

Present results showed that transginic maize and soybean crops are not cultivated in the studied areas of Ethiopia. These were identified by specific primers 35S and specific for endgnous genes of maize and soybean. The amplicons of the specific sequences detected are used to build the different GMOs and regulate expression of transgenes, such as promoter 35S and the terminator NOS [20]. Furthermore, PCR is a useful machine in detecting the transgenic plants [21].

From the time when first generation of GMOs seed were released for plantation in 1995 and their products entered the food and feed markets of various countries in 1996 [22], there have been reports questioning the safety of these organisms and their products when used as food or feed. This brought about an active debate and controversy on possible risks that might be caused by such crops and their products to health [23] and environment [24]. As a result two main methods for the identification of GM food and feed have been reported in the literatures; PCR and Enzyme-Linked Immunosorbent Assay (ELISA). PCR is the most accepted technique used worldwide; it showed consistent results when using specific primers for the detection of the regulatory sequence or structural gene in the inserted gene fragment [25]. The designed primers must possess some specific characteristics and can be used for GM product screening and product-specificity detection. Thus, it might be convenient to consider these methods for the detection and identification of GMOs. This would represent a new field of diagnostics in which a great deal of development has already been accomplished.

## 7. CONCLUSION

Finally, the soybean and maize has been tested for the presence or absence transgenes using thermal cyclere PCR, fortnutelly there is no GMO crops in the studied area. The protocol developed can be used to meet the regulatory obligations and for the labeling of GM plants and their derivatives for future in Ethiopia. The PCR method described in the study is an efficient and sensitive tool, which can be applied for routine screening/testing of herbicide resistant GM soybean and GM maize from conventional/non-GM soybean or maize.

## 8. ACKNOWLEDGMENTS

Ethiopian Enviroment and Forest Research Institute appreciates highly Ethiopian Enviroment, Forest and Climate Change Commision for supporting this study by purchasing molculare biology reagents and equipments which are used for GMO detection.

## Refferance

1. Antony Ceasar S, Ignacimuthu S. Genetic engineer-ing of crop plants for fungal resistance: role of anti-fungal genes. Biotechnol Lett 2012; 34:995-1002; PMID:22350290; http://dx.doi.org/10.1007/s10529-012-0871-1.2.

2. Sheludko YV. Recent advances in plant biotechnol-ogy and genetic engineering for production of sec-ondary metabolites. Tsitol Genet 2010; 44:65-75; PMID:2 0201415.3.

3. Thao NP, Phan Tran LS. Potentials toward genetic engineering of drought-tolerant soybean. Crit Rev Biotechnol 2011; PMID:22181694; http://dx.doi.org/10.3109/07388551.2011.643463.4.

4. Yang S, Vanderbeld B, Wan J, Huang Y. Narrowing down the targets: towards successful genetic engineer-ing of drought-tolerant crops. Mol Plant 2010; 3:469-90; PMID:20507936; http://dx.doi.org/10.1093/mp/ssq016.5.

5. Wally O, Punja ZK. Genetic engineering for increasing fungal and bacterial disease resistance in crop plants. GM Crops 2010; 1:199-206; PMID:21844674; http://dx.doi.org/10.4161/gmcr.1.4.1

6. Kumar V, Parkhi V, Kenerley CM, Rathore KS. Defense-related gene expression and enzyme activities in transgenic cotton plants expressing an endochitinase gene from Trichoderma virens in response to interac-tion with Rhizoctonia solani. Planta 2009; 230:277-91; PMID:19444464; http://dx.doi.org/10.1007/s00425-009-0937-z.7.

7. Parkhi V, Kumar V, Campbell LM, Bell AA, Shah J, Rathore KS. Resistance against various fungal patho-gens and reniform nematode in transgenic cotton plants expressing Arabidopsis NPR1. Transgenic Res 2010; 19:959-75; PMID:20151323; http://dx.doi.org/10.1007/s11248-010-9374-9

8. ISAAA: International Service for the Acquisition of Agri-biotech Application. 2011. Available from: URL: http://www.isaaa.org

9. Margarit E, Reggiardo MI, Vallejos H, Permingeat H R. Detection of BT transgenic maize in foodstuffs. Food Res. Int. 2006;39:250–255.

10. Matsouka T, Kuribara H, Akiyama H, Hirohito M, Goda Y, Kusakabe Y, Isshiki K, Yoyoda M, Hino A. A multiplex PCR method of detecting recombinant DNAs from five lines of genetically modified maize. J.food Hyg. Soc. Japan. 2000;42:24–32.

11. Kakihara y, Matsufuji H, Chino M, Yamagata K. Detection of recombinant DNA of genetically modified GM soybeans in heat-treated GM soybeans and commercial natto. Food Control. 2007;18:1289–1294.

12. Deisingh AK, Badrie N. Detection approaches for genetically modified organisms in foods. FoodResearch International. 2005;38:639–649.

13. Yamaguchi H, Sasaki K, Umetsu H, Kamada H. Two detection methods of genetically modified maize and the state of its import into Japan. Food Control. 2003;14:201–206.

14. Holst-Jensen A. Testing for genetically modified organisms (GMOs): past, present and future perspectives. Biotechnology advances. 2009;27:1071–1082.

15. Miraglia M, Berdal KG, Brera C, Corbisier P, Holst-Jenen A, Kok EJ, Marvin HJP, Schimmel J, Rentsch J, Rie JPPF, Zagon J. Detection and traceability of genetically modified organisms in the food production chain. Food and Chemical Toxicology. 2004;42:1157–1180.

16. Germini A, Zanetti A, Salati C, Rossi S, Forre C, Schmid S, Marchelli R. Development of a seventarget multiplex PCR for the simultaneous detection of transgenic soybean and maize in feeds and foods. JAgric. Food chem. 2004;52:3275–3280.

17. Rabiei, M, Mehdizadeh M, Alebouyeh M, Rastegar H. Screening of genetically modified organisms and specific detection of Bt-11 maize in foodstuffs; April 30th - May 1st; Tehran, Iran: Proceedings of the 2nd International Congress of FoodHygiene; 2011.

18. Meyer R, Chardonnens F, Hubner P & Luthy J, Polymerase chain reaction (PCR) in the quality and safety assurance of food: Detection of soya in processed meat products, ZLebensm Unters Forsch, 203 (1996) 339–344.

19. Ehlers V B, Strauch E, Goltz M, Kubsch D, Wagner H et al, Nachweis gentechnischer Veranderungen in Mais mittels PCR, Bundesgesundheitsblatt, 4 (1997) 118–121.

20. Vijayakumar, K.R., A. Martin, L.R. Gowda and V. Prakash, 2009. Detection of genetically modified soya and maize: Impact of heat processing. Food Chem., 117: 514–521.

21. Cankar, K., V. Chauvensy-Ancel, M.N. Fortabat, K. Gruden, A. Kobilinsky, J. Zel and Y. Bertheau, 2008. Detection of nonauthorized genetically modified organisms using differential quantitative polymerase chain reaction: Application to 35S in maize. Anal. Biochem., 376: 189–199.

22. Paarlberg, P., 2006. Are Genetically Modified (GM) crops a commercial risk for Africa. Int. J. Technol. Globalisation, 2: 81–92.

23. Al-Jebreen, D.H., 2010. Perceptions and attitudes of riyadh university students towards products derived from genetically modified crops in Saudi Arabia. Pak. J. Biol. Sci., 13: 28–33.

24. Kawata, M., K. Murakami and T. Ishikawa, 2009. Dispersal and persistence of genetically modified oilseed rape around Japanese harbors. Environ. Sci. Pollut. Res. Int., 16: 120–126.

25. James, D., A.M. Schmid, E. Wall, M. Green and S. Masri, 2003. Reliable detection and identification of genetically modified maize, soybean and canola by multiplex PCR analysis. J. Agric. Food Chem., 51: 5829–5834.

